# A global map of receptor-binding protein compatibility for the programmable design of Klebsiella and Acinetobacter phages

**DOI:** 10.64898/2026.02.20.706991

**Authors:** András Asbóth, Tamás Stirling, Orsolya Méhi, Gábor Apjok, Victor Klein De Sousa, Nicholas M I Taylor, Hiba Hadj Mehdi, Balázs Papp, Eszter Ari, Bálint Kintses

## Abstract

The narrow host range of bacteriophages limits their application against genetically diverse bacteria, motivating rational host-range modification. Progress in programmable engineering depends on establishing design principles that determine the compatibility of possible host-recognition modules with phage scaffolds. Here, we establish a genomics-guided framework that systematically identifies compatibility-determining adapter domains in receptor-binding proteins for the rational engineering of phages to target clinically relevant pathogen populations. Applying this approach to 1,270 phage genomes infecting *Acinetobacter baumannii* and *Klebsiella pneumoniae*, we show that viral diversity is highly structured: 60% of the 2,313 receptor-binding proteins group into only 19 major clusters sharing conserved N-terminal compatibility adapters. The structurally most conserved adapters in Autographivirales are associated with diverse capsule-degrading depolymerases. Known host-specificities within single-adapter clusters target capsule types that represent up to 29% of *A. baumannii* and 44% of *K. pneumoniae* carbapenem-resistant populations. Overall, we define a global repertoire of modular host-recognition components for programmable configuration of phage therapeutics.

## Background

Antibiotic resistance is projected to become the leading cause of death by 2050, underscoring the urgent need for alternative treatments.^1^ Among the most concerning trends is the rise of pan-resistant infections – cases in which no effective antibiotics remain.^2^ Phage therapy, which uses viruses to selectively kill bacteria, has emerged as a promising alternative.^3,4^ While growing clinical evidence supports its use in compassionate use for antibiotic-resistant infections,^5,6,7,8,9^, broad clinical translation remains limited.^10,11^ A major obstacle is the narrow host range of most bacteriophages, which restricts their utility against genetically diverse pathogens.^12,13^ Even treating a single bacterial species often requires a large and diverse phage library isolated from nature.^14,15^

This challenge is especially pronounced for two ESKAPE pathogens: *Acinetobacter baumannii* and *Klebsiella pneumoniae*. Carbapenem-resistant strains of both species top-ranked among the WHO’s critical pathogens list due to their high mortality rates and limited treatment options.^2^ Their thick, variable polysaccharide capsules impede phage binding to the cell surface, demanding highly tailored phage libraries to achieve therapeutic coverage^14,16,17^. With 237 and 187 known capsule locus variants in *A. baumannii* and *K. pneumoniae*, respectively, ^18,19,20^ targeting even only the most common carbapenem-resistant capsule types requires extensive diversity among phage candidates.^14^ At present, building such libraries necessitates isolating and characterizing natural phages and conducting separate manufacturing, pre-clinical, and clinical development for each agent at a scale that renders phage therapy financially non-viable.^21^ To significantly reduce these development costs and time, one solution could be to engineer already validated phages to retarget them to new hosts without full-scale redevelopment for each new variant.

Phage engineering studies have demonstrated that receptor-binding proteins (RBPs) – the major host recognition component of phages — can transfer host specificity between phages when swapped.^22,23,24,25,26,27^ This is made possible by the modular structure of RBPs ^23,25,28,29^, typically composed of three parts: i) a C-terminal receptor-binding domain that recognizes host bacterial cell surface receptors, such as polysaccharides, proteins, or lipopolysaccharides, ii) an optional upstream depolymerase domain, which degrades bacterial capsules, and thereby defines capsule-type specificity, and iii) an N-terminal adapter domain that anchors the RBP to the phage particle and ensures compatibility with the viral scaffold^23,25,30^ (Figure 1A). In nature, RBPs are transferred between phages as a full unit or in parts—sometimes excluding the adapter, when it’s incompatible with the recipient phage.^28,31,32,33^ Similarly, phage engineering follows two main strategies: i) “plug-and-play,” where a fully compatible RBP is available and can be inserted in the phage genome without alteration, or ii) custom fusion, where engineering of an RBP is required to merge a recipient phage’s adapter with the incoming functional domains to make it compatible with the phage scaffold.^24,25,27,34,35^ Although plug-and-play RBP integration is an efficient strategy, success depends on identifying RBPs that are structurally and functionally compatible with the phage scaffold. Currently, however, the natural diversity of such compatible RBPs remains poorly mapped. Expanding this knowledge is critical to scale programmable phage design from successful examples to full-scale coverage of pathogen populations.

**Figure 1.**
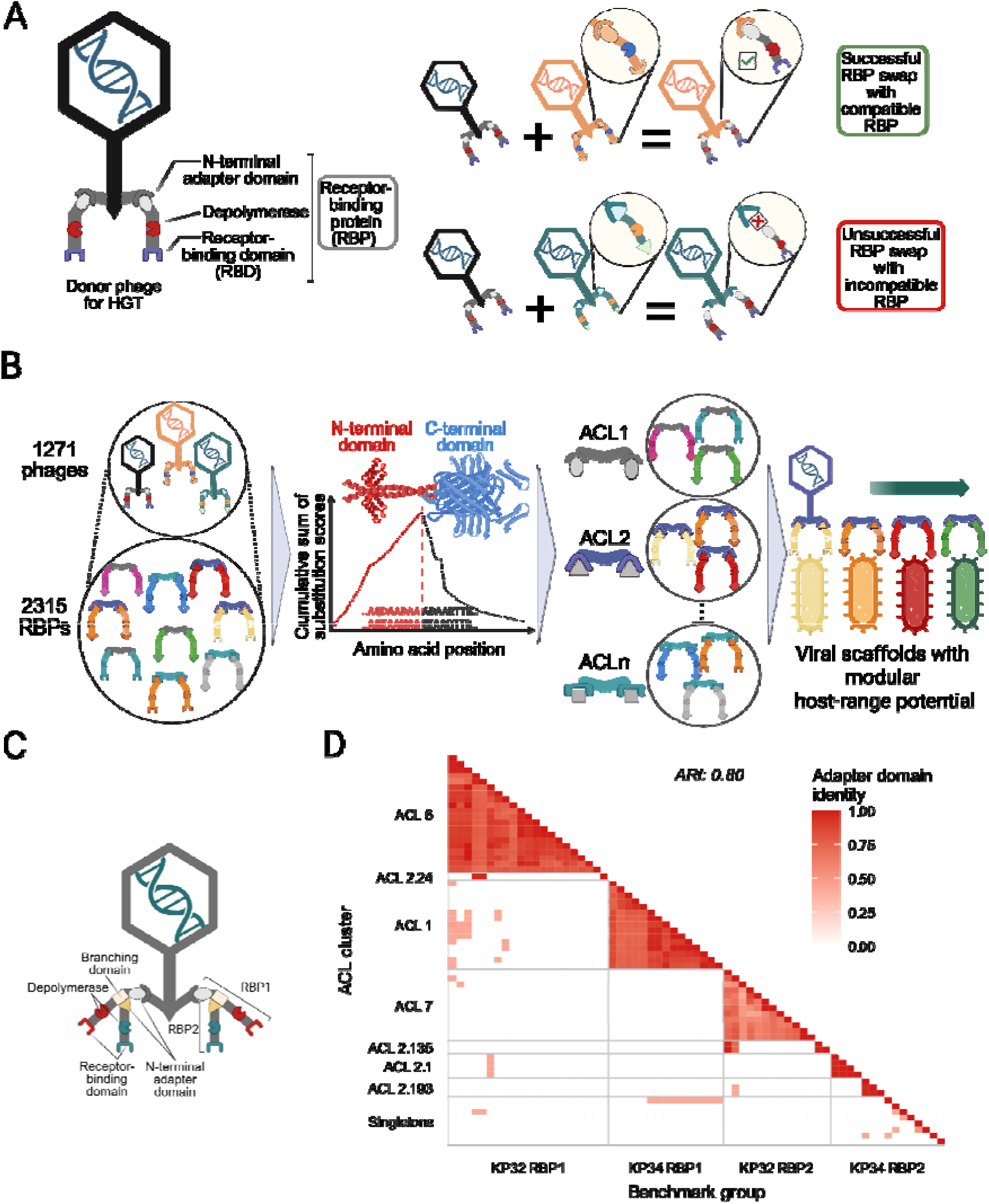
Large-scale detection and clustering of RBPs’ N-terminal adapter domains. **A)** Schematic representation of the modular architecture of a phage RBP and illustration of two RBP swap scenarios with compatible and incompatible RBPs, respectively. **B)** Schematic representation of the comparative genomics pipeline involving i) identification of RBPs from phage genomes, ii) prediction of N-terminal adapter domains based on amino acid conservation, iii) grouping of adapter domains into adapter clusters (ACLs), and iv) matching RBPs with phage host specificities to create interchangeable phage libraries **C)** Schematic model (based on Latka et al., 2019) of the anchor-branched RPB system of a KP32 phage with dual receptor specificity, possessing two RBPs. The second RBP attaches to the first via a branched configuration. **D)** Clustering RBPs of 35 Klebsiella phage genomes from a manually curated dataset (Latka et al., 2019) (Table S1) to validate the TAILOR pipeline. Three benchmark groups – KP32 RBP1 (corresponding to ACL 6), KP32 RBP2 (corresponding to ACL 7), and KP34 RBP1 (corresponding to ACL 1) – out of the four were identified by high accuracy (ARI = 0.8) by our pipeline.

Here, we establish a genome-sequence-based framework to map the integratable repertoire of RBPs in *A. baumannii* and *K. pneumoniae* phages. Using TAILOR (Tool for Adapter-domain Identification and Linking Of RBPs (https://github.com/sbthandras/tailor)), we automatically identify adapter domains that anchor RBPs to phage scaffolds. Analysis of 2,313 RBPs from Acinetobacter and Klebsiella phages shows that most RBPs are anchored to the phage particle through a limited set of highly similar adapter domains. Several Autographivirales genera emerge as particularly promising engineering scaffolds, combining conserved adapter-mediated anchoring with depolymerase enzymes targeting a wide range of clinically relevant bacterial capsules. The resulting global RBP repertoire also reveals key evolutionary patterns of RBP exchange among phages.

## Results

### Mapping receptor-binding protein compatibility

Depolymerase enzymes in RBPs often undergo horizontal gene transfer. This creates a unique genomic pattern: a conserved N-terminal region paired with a highly variable, recombination-prone C-terminal region.^32^ Our comparative genomics pipeline, TAILOR, is designed to exploit this pattern. Using full-length amino acid sequences of RBPs, TAILOR detects the boundary between the conserved N-terminal domain and the variable downstream region. It then clusters RBPs based on the sequence identity of their conserved N-terminal domains (See Methods and Figure 1B). To feed into TAILOR, we retrieved all publicly available *A. baumannii* and *K. pneumoniae* phage genomes as of 2025 January, yielding a dataset of 1,270 phage genomes containing 2,313 RBPs (see Methods). We then performed pairwise global alignments of all unique RBP sequences to detect a shift in amino acid sequence identity. Based on these, we grouped RBPs with highly similar adapter domain sequences into adapter clusters, providing the basis for subsequent comparative analysis (Figure 1B, Figure S1, Methods, and Tables S1 and S2).

We validated our clustering strategy against a benchmark subset of RBPs from 36 Klebsiella phage genomes of the genera Przondovirus and Drulisvirus – commonly referred to as KP32 and KP34 phages – that were characterized in a previous study (Table S3).^32^ Most of these genomes encode two RBPs with manually annotated N-terminal adapter domains.^32^

The first RBP attaches directly to the phage scaffold through a highly conserved adapter domain longer than 100 amino acids, while the second RBP connects to the first one with a shorter, less conserved adapter (Figure 1C). In this way, these RBPs cluster into four distinct adapter-domain groups (KP32-RBP1, KP32-RBP2, KP34-RBP1, and KP34-RBP2), each differing in domain length and conservation.^32^ Our method accurately clustered three out of the four benchmark adapter groups, achieving an Adjusted Rand Index (ARI) of 0.8^36^, where an ARI of 1 represents perfect clustering agreement (Figure 1D, Table S4). The KP34-RBP2 group was not clustered by our method, most likely because its adapter domains are exceptionally short (only 7 amino acids long) and contain only ∼4 conserved positions on average, below the sensitivity of TAILOR (Figure 1D, see Methods). Finally, comparison of the predicted adapter lengths to those of the manually curated adapters showed strong agreement, with a median difference of 2.9% of the adapter length between predicted and observed domain boundaries, supporting the accuracy of TAILOR’s N-terminal adapter identification (Table S3). Applying this framework to the full dataset of 2,313 RBPs revealed a highly structured landscape. Sequence-based clustering revealed 374 distinct adapter clusters (median predicted adapter length of 158 amino acids), of which 172 are non-singletons (Figure 2A, Figure S2, Tables S1 and S5). Strikingly, the distribution of cluster sizes is heavily skewed: 58% of all RBPs fall into just 19 large adapter clusters, each containing at least 30 members (Table 1, Figure 2A, Methods). Within these large clusters, adapter domains share an average of 79% amino acid sequence identity and are typically restricted to phages belonging to the same genus (Figure 2A, B). Occasionally, cross-genus conservation was also observed across genera of the same phage family (Figure 2A), albeit with a lower average sequence identity compared to adapter domain pairs from the same genus (Figure 2B, Supplementary File 1). Overall, large adapter clusters are common across multiple phage families (Figure 2A), including representatives from all three major phage morphotypes — Siphophage, Myophage, and Podophage. Finally, to assess whether the observed sequence identities within adapter groups are potentially sufficient to enable RBP exchange, we collected documented cases of successful RBP swaps from the literature (Table S6). We identified 13 such cases, most often involving phages from different genera, predominantly from the order Autographivirales. We analyzed the sequence identity of the swapped adapter domains and found an average amino acid identity of 76%, with the lowest successful swap occurring at 61% identity. These findings support our hypothesis that sequence identities within observed adapter clusters can be sufficient to enable functional exchange of RBPs among phages within the same cluster.

**Figure 2.**
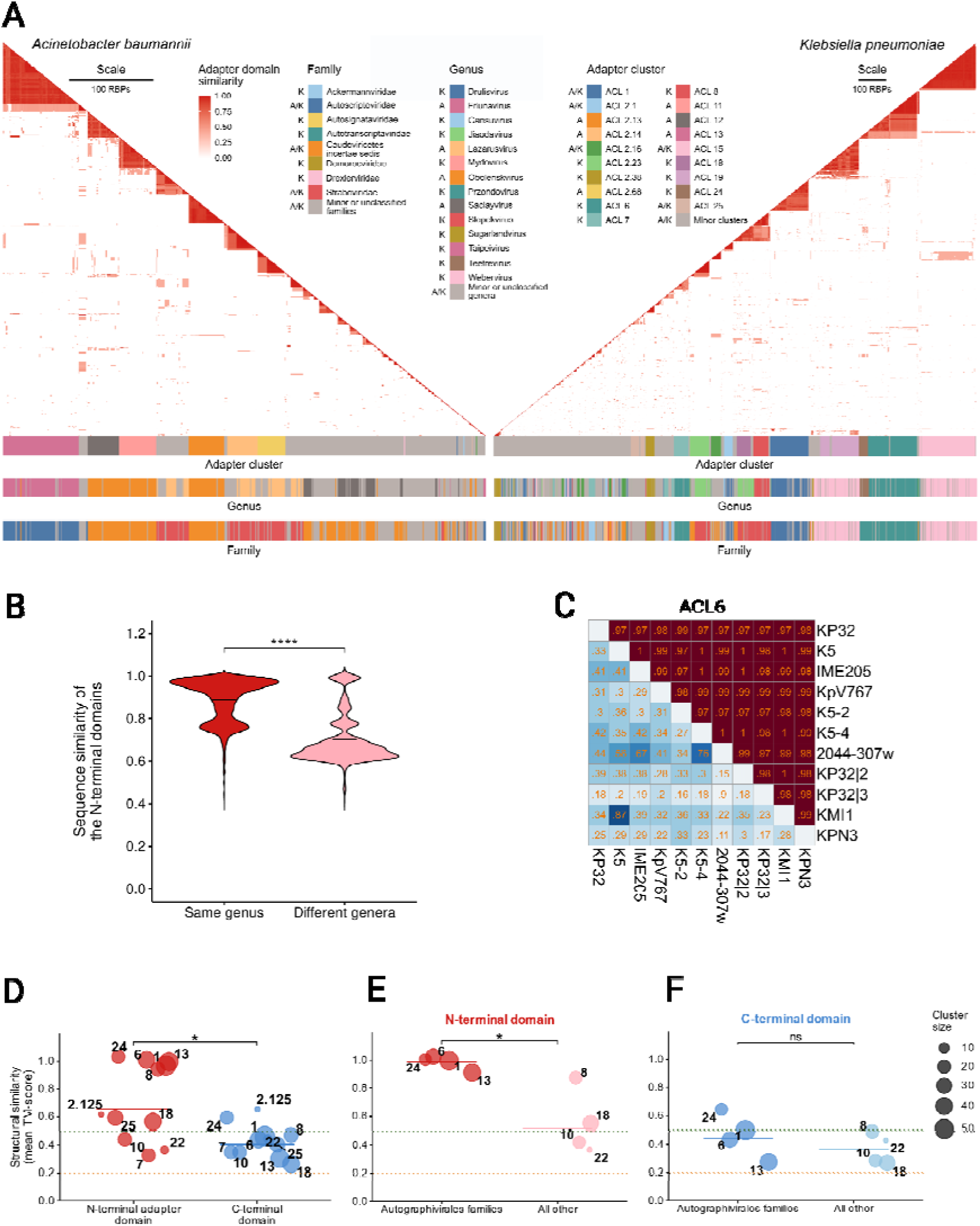
Structural conservation of N-terminal adapter domains within adapter clusters. **A)** Clustering of the 2,313 RBPs using TAILOR identified 172 distinct non-singleton adapter clusters. Red intensity indicates adapter domain sequence similarity, darker shades representing higher similarity. Clusters are labeled by number and color-coded by phage taxonomy (genus and family). Host species are indicated by A (*A. baumannii*), K (*K. pneumoniae*) and A|K, for mixed clusters (*N* = 574 and 1,739 RBPs for *A. baumannii* and *K. pneumoniae*, respectively). **B)** N-terminal sequence similarity within clusters is higher for phages from the same genus than from different genera (mean 0.89 *vs.* 0.68; *N* = 99,964 and 3,564 comparisons). Only RBPs from the 19 largest clusters were analyzed (Table 1, Figure 2A); similarities < 0.4 were excluded. Two-tailed Wilcoxon rank-sum test, *p* < 2.2 × 10lJ¹lJ. **C)** Heatmap of pairwise TM-score structural similarities of N-terminal adapter domains (upper right, red) and C-terminal domains (lower left, blue) within ACL 6. TM-scores are shown in each cell, phage names on the *x* and *y* axes (N = 11). **D)** Within clusters, N-terminal domains show greater structural similarity than C-terminal domains (Wilcoxon rank sum test, *p* = 0.061; *N* = 11 clusters). Each datapoint represents the average pairwise TM-score within an adapter cluster. **E)** Among adapters >100 amino acids, those from Autographivirales families exhibit higher within-cluster similarity (mean TM-score 0.981) than those from other families (*p* = 0.0061; *N* = 4 per group). **F)** In contrast, C-terminal domains from Autographivirales families do not show higher similarity (*p* = 0.93; *N* = 4 per group). In D–F, points represent mean TM-scores per cluster; dot size reflects cluster size. TM-score thresholds: 0.2 (random similarity), 0.5 (shared fold), 1 (identical structures). Orange and green dotted lines mark TM-score thresholds of 0.2 and 0.5, respectively.

**Table 1.** Large adapter clusters with at least 30 RBPs. Abbreviations: A: *A. baumannii*, K: *K. pneumoniae*, A|K: clusters containing both *A. baumannii* and *K. pneumoniae* phages, Drex: Drexlerviridae, Autotr: Autotranscriptaviridae, Autosc: Autoscriptoviridae, Strab: Straboviridae, Chima: Chimalliviridae, Deme: Demerecviridae. Adapter pident is the median percentage identity of all pairwise comparisons of adapter domain sequences within unique RBPs of an adapter cluster (see Methods). The value for ACL 2.16 is missing because it contains only one unique RBP. The “Enzyme clusters” column contains the number of detected enzyme clusters within each adapter cluster. The “CPS types” column contains the number of *A. baumannii* and/or *K. pneumoniae* CPS types that the phages within the cluster may infect.

| Adapter cluster | Host | Phage family | Nr. of phages | Nr. of RBPs | Adapter pident | Enzyme clusters | CPS types |
| --- | --- | --- | --- | --- | --- | --- | --- |
| ACL 15 | A\|K | Drex | 195 | 203 | 0.944 | 2 | 2 |
| ACL 6 | K | Autotr | 165 | 173 | 0.864 | 18 | 11 |
| ACL 1 | A\|K | Autosc | 129 | 139 | 0.787 | 14 | 6 |
| ACL 19 | K | Drex | 123 | 131 | 0.882 | 14 | 0 |
| ACL 13 | A | Autosc | 77 | 90 | 0.909 | 36 | 17 |
| ACL 2.23 | K | Strab | 53 | 70 | 0.615 | 0 | 0 |
| ACL 8 | K | Strab | 51 | 54 | 0.940 | 0 | 0 |
| ACL 7 | K | Autotr | 48 | 53 | 0.610 | 18 | 4 |
| ACL 18 | K | Strab | 41 | 50 | 0.888 | 2 | 0 |
| ACL 25 | A\|K | Chima | 35 | 47 | 0.443 | 21 | 0 |
| ACL 11 | A | unspecified | 41 | 44 | 0.897 | 0 | 0 |
| ACL 2.13 | A | unspecified | 40 | 42 | 0.889 | 0 | 0 |
| ACL 12 | A | unspecified | 35 | 37 | 0.843 | 16 | 1 |
| ACL 2.16 | A\|K | unspecified | 37 | 37 |  | 0 | 0 |
| ACL 2.14 | A | Strab | 28 | 36 | 0.655 | 1 | 1 |
| ACL 2.38 | K | Deme\|Strab | 32 | 33 | 0.737 | 2 | 0 |
| ACL 2.68 | A | Strab | 26 | 33 | 0.482 | 0 | 0 |
| ACL 24 | K | Autotr | 33 | 33 | 0.893 | 3 | 0 |
| ACL 2.1 | A\|K | Autosc | 28 | 32 | 0.982 | 2 | 1 |

### Adapters with exceptional structural conservation

Next, we evaluated whether conserved adapter sequences translate into similarly conserved structures. Therefore, we generated computationally predicted 3D structures for a representative set of 105 RBPs from 28 adapter clusters. First, by visual inspection, we compared adapter boundaries predicted by TAILOR with those in the 3D models to further assess the accuracy of our predicted adapter domains (see Methods). Consistent with TAILOR’s limited sensitivity to extremely short adapters (Figure 1D), adapter domains shorter than 15 amino acids were generally not detected (Figure S3). The 3D structures also revealed that the boundaries of adapters longer than 200 amino acids are often artefacts arising from the presence of additional conserved domains downstream of the N-terminal adapter domain (Figure S3). For the remaining adapter clusters, the two methods generally agreed on the N-terminal domain boundaries, with the predicted and observed boundaries differing by a median of 25% of the observed adapter length (Figure S3, Table S7).

Then, we calculated Template Modeling scores (TM-scores) for each RBP structure pair – a metric that assesses the extent to which two protein structures superimpose (see Methods). A separate TM-score was calculated for the N-terminal adapter and for the remainder of the protein up to the C-terminus (Figure 2C, Figure S4, Table S8 and S9). As expected, the N-terminal adapter domains displayed significantly higher structural similarity than the variable C-terminal domains (Figure 2D, Figure S4, Table S10), supporting the idea that conserved sequences in adapter domain clusters reflect conserved structure. However, structural conservation levels varied across adapter clusters, and a striking pattern emerged. In families belonging to the order Autographivirales (Autonotataviridae, Autoscriptoviridae, Autosignataviridae, and Autotranscriptaviridae), adapter domains longer than 100 amino acids displayed near-perfect within-cluster structural similarity, which is significantly higher than that of adapters from other families (Figure 2E, Figure S4, Table S11). Additionally, despite their highly similar adapters, the C-terminal domains of RBPs from Autographivirales families show similarly high structural divergence compared to that of RBPs from other families (Figure 2F, Table S12). This indicates that these RBPs are functionally diverse despite their uniform adapters. Notably, Autographivirales adapter domains shorter than 100 amino acids (typically 20-36 amino acids), such as those in KP32-like phages that form ACL7, exhibit lower structural similarity (Figure S4). These adapters connect to an upstream RBP rather than directly to the viral scaffold^32^ (Figure 1C). This suggests that RBPs that rely on such short adapters are likely swappable only in tandem with their upstream RBP, functioning as a combined structural unit rather than as independent modules.

In sum, we found that Autographivirales phages stand out with highly conserved adapter domains, making them particularly well-suited for the configuration through complete RBP swaps.

### Conserved viral adapters anchor diverse enzymatic repertoires

We next examined the diversity of depolymerase enzymes and their capsule-specificities within adapter clusters to estimate the extent of possible host-range modifications for a specific viral scaffold. To do this, we first annotated depolymerases within the RBPs using previously established tools (see Methods). Then, to assign specificities to them, we assembled a literature-curated dataset of RBPs with experimentally determined or predicted capsule specificities (Table S13, see Methods). The dataset included 32 RBPs from *A. baumannii* phages and 215 from *K. pneumoniae* phages with 23 and 67 known capsule targets, respectively. Most depolymerases originated from phages encoding a single enzyme (typically Autographivirales), enabling unambiguous specificity assignment, whereas phages carrying multiple depolymerases were less informative because individual enzymes could not be reliably linked to capsule targets.^37^ We then grouped the detected enzymes in the RBPs into 220 enzyme clusters using a 90% amino acid sequence identity threshold and annotated each cluster with the known specificity of any RBP it contained. Finally, we defined the functional repertoire of each adapter cluster by quantifying the number of enzymatic clusters with known versus unknown capsule targets within each adapter cluster.

Among the 2,313 RBPs, only 31% (713 RBPs) contained an annotated depolymerase (Figure 3A). Out of these, 245 RBPs could be assigned a capsule specificity (Figure 3A, Table S14). This leaves many RBPs with unexplored functional potential, likely encoding either undetected depolymerases or RBPs with functions other than capsule degradation. ^38^

**Figure 3.**
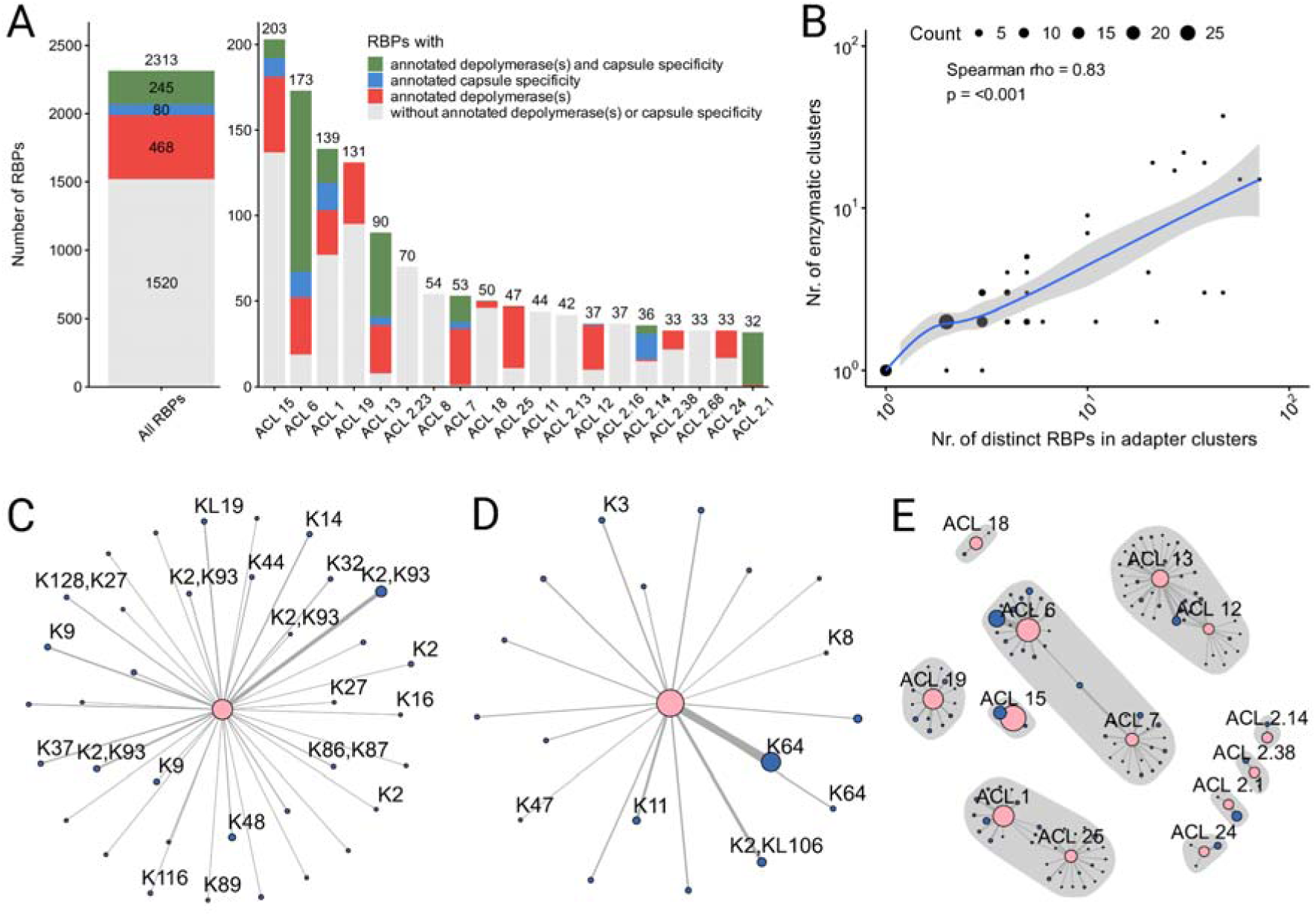
Enzymatic diversity within adapter clusters. **A)** Number of RBPs in adapter clusters, grouped by available depolymerase and specificity annotations. The left panel shows all clusters combined, while the right panel shows the top 19 clusters (Table 1). Column heights, and numbers above each column show the total number of RBPs assigned to each adapter cluster. The stacked bar segments colored green, blue, red and grey represent the number of RBPs with annotated depolymerase(s) and capsule specificity, annotated capsule specificity only, annotated depolymerases only, and without any annotated depolymerase(s) or capsule specificity, respectively (see Methods). **C)** The number of distinct enzyme clusters within an adapter cluster scales linearly with adapter cluster size. Each point belongs to one or more adapter clusters. The size of each point reflects the number of overlapping adapter clusters in that location. Larger adapter clusters tend to have more enzyme clusters (Spearman correlation, *N* = 73, *rho* = 0.63, *p* < 0.01) with no observed plateau. **C)** and **D)** Enzyme clusters and capsule specificities associated with RPBs in the largest *A. baumannii* adapter cluster (ACL13) and *K. pneumoniae* adapter cluster (ACL 6), respectively. The pink node indicates the adapter cluster, while the blue nodes indicate enzyme clusters within that adapter cluster. Labels for enzyme clusters indicate associated capsule specificities (see Methods). **E)** Network of linked adapter and enzyme clusters for RBPs belonging to the 19 largest adapter clusters (Table 1). Six adapter clusters containing RBPs without any annotated depolymerase(s) or capsule specificity (grey bars in panel A) are not shown. Linked adapter and enzyme clusters define a component, isolated components are highlighted in grey.

Interestingly, the number of distinct enzyme clusters within an adapter cluster scaled linearly with adapter cluster size without any sign of saturation (Figure 3B, Table S15). The largest adapter clusters for *A. baumanni* phages (ACL 13) and *K. pneumoniae* phages (ACL 6) were also among the most diverse, containing 36 and 18 enzyme clusters, respectively (Figure 3C-D, Tables S16 and S17). Many of these have known specificities, highlighting their potential to serve as a basis for programmable phage specificity. Finally, across the 19 large adapter clusters, we observed that each cluster tends to have a unique depolymerase repertoire: 91% of enzymatic clusters were associated with a single adapter cluster, whereas the remaining were shared across two adapter clusters (Figure 3E, Tables S18 and S19). These overlaps likely reflect recent recombination events between adapters and depolymerases that occurred during the horizontal transfer of these depolymerase genes.

### A limited set of adapter clusters covers a major fraction of global carbapenem-resistant lineages

Having shown that RBPs often share highly conserved adapter domains while harboring diverse enzymatic domains, we asked whether RBPs from a single adapter cluster could target a broad set of clinical strains. If true, this would imply that a small number of phage scaffolds equipped with diverse, compatible RBPs might be sufficient to treat a wide range of infections. To test this hypothesis, we first estimated the prevalence of capsule types among carbapenem-resistant *A. baumannii* and *K. pneumoniae*. To this end, we downloaded and processed all high-quality, human-derived, publicly available *assembled genomes of A. baumannii and K. pneumoniae* collected between 2018 and 2023 that carried one or more carbapenem resistance determinants. To minimize sampling bias inherent in a public database, we stratified both datasets by country-level population size using available metadata, following an established pipeline^14^ (see Methods). Overall, from the available 30,827 *A. baumannii* and 55,742 and *K. pneumoniae* genomes, quality filtering, removal of carbapenemase-non-producing genomes, and geographic stratification yielded 2,946 *A. baumannii* and 2,959 *K. pneumoniae* genomes from the last 6 years, spanning 91 countries across 18 world regions out of 22, representing the current populations of these pathogens (Table S20). Then, we matched the capsule types of these genomes to the capsule specificities of the RBPs in the adapter clusters. For carbapenem-resistant *A. baumannii*, RBPs with known capsule specificities cover 44.9% of the global pathogen’s population (Figure 4A). The largest share of this coverage is provided by RBPs from ACL 13, with 10 known capsule-specificities. These RBPs belong to phages in the genera Friunavirus and Obolenskvirus within the order Autographivirales (Figure 4A, Table S21). Importantly, Friunavirus RBPs in ACL 13 target globally dominant capsule types, such as K2, K9, and K14, which together account for 27% of the investigated carbapenem-resistant *A. baumannii* population (Figure 4B, Table S22) while the other 7 CPS types in ACL 13 (K32, K16, K116, K37, K48, K44, K127) account for an extra 2%. Additionally, Friunaviruses have been used for therapy^7,39^, and therefore, they can be considered as a phage scaffold with therapeutic potential. Notably, significant gaps remain: the canonical K3 capsule (the second most prevalent among carbapenem-resistant *A. baumannii*) lacks a known Friunavirus RBP. The only reported K3-specific Friunavirus targets a minor K3 variant.^40^

**Figure 4.**
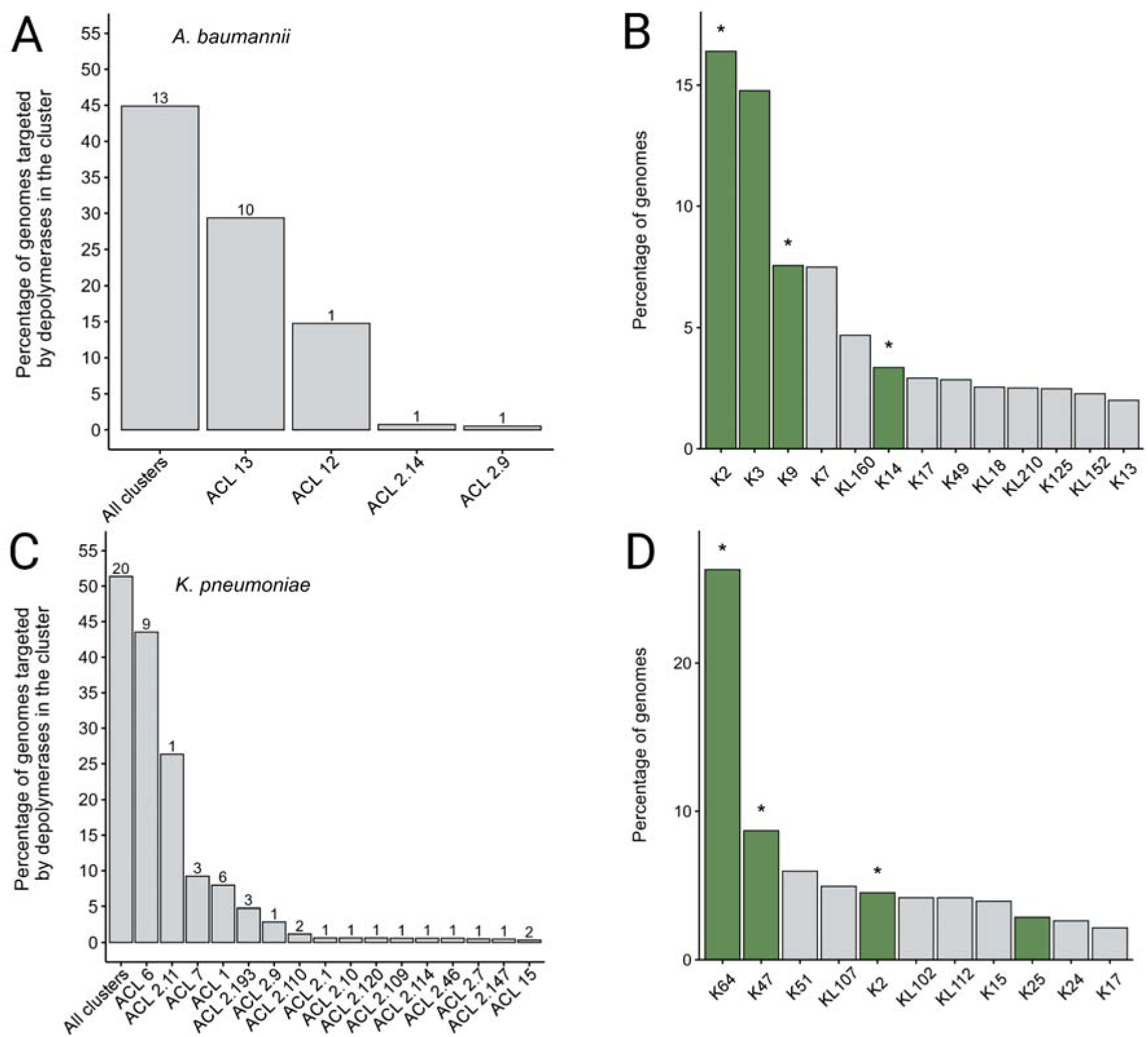
A large proportion of the global carbapenem-resistant lineages can be covered by RBPs belonging to a limited set of adapter clusters. Coverage of the global *A. baumannii* (A) and *K. pneumoniae* (C) population by RBPs associated with known capsule polysaccharide (CPS) specificities. The *y*-axis displays the percentage of *A. baumannii* (A) or *K. pneumoniae* (C) genomes belonging to CPS types associated with RBPs in a given adapter cluster or in all adapter clusters (first bar). Only large adapter clusters are shown that contain RBPs with known CPS specificities (see Table 1), with the exception of the first bar, where all clusters were considered. The numbers above the bars show the number of distinct CPS specificities associated with RBPs in a given adapter cluster or all adapter clusters. Relative global prevalence of the most prevalent *A. baumannii* (**B**) and *K. pneumoniae* (**D**) CPS types that together account for 70% of worldwide genomes (Table S22, *N* = 2,117 and 2,080 for *A. baumannii* and *K. pneumoniae* genomes, respectively). For capsule types highlighted with green we identified phages with corresponding capsule specificities. Stars mark capsule types associated with RBPs from the ACL 13 (*A. baumannii*) and ACL 6 (*K. pneumoniae*) clusters, respectively.

For carbapenem-resistant *K. pneumoniae,* 51.4% of the population is covered by RBPs with known specificities (Figure 4C). This coverage is driven by RBPs from the largest adapter clusters (ACL 1, 6, and 7), which belong to phages from the order Autographivirales (Figure 4C, Table S21). ACL 6 contributes the most to the coverage by harbouring RBPs that target 9 different CPS types, including globally dominant capsule types, such as K64, K47 and K2, which together account for 39.5% of the global population (Figure 4D, Table S22) while the other 6 CPS types in ACL 6 (KL106, K20, K30, K3, K57, K8) account for an extra 4%. Despite this breadth, key types remain uncovered; no RBP in these clusters targets KL107, one of the most prevalent capsules (4.9%), underscoring the need for further discovery (Figure 4D). RBPs in the ACL 6 cluster belong to phages from the genus Przondovirus within the order Autographivirales, which have been used in several clinical case studies ^7,41,42^, reinforcing the clinical relevance of these phages as a therapeutic scaffold.

We also observed that the most prevalent capsule types are often targeted by multiple viral operational taxonomic units (vOTU, that is <95% sequence identity), usually through identical or highly similar RBPs. In particular, 94 distinct *Klebsiella* vOTUs share an RBP with very high sequence identity along the entire length, including the adapter that belongs to ACL 6 (Figure S5). These RBPs are specific to K64, the most common capsule type among carbapenem-resistant *K. pneumoniae* (Figure S5, Table S23). Similar patterns were seen for K24 and K1 specific Klebsiella phage RBPs found in adapter ACL 15 and 2.1, respectively, and a parallel trend is evident in *A. baumanii* among the K2 capsule-specific RBP in ACL 13. These observations suggest either that RBPs are relatively conserved compared to other regions of phage genomes and thus remain identical while the rest of the genome diversifies, or that horizontal transfer of RBPs targeting highly prevalent serotypes is positively selected, leading to their widespread distribution across diverse phage genomes.

## Discussion

Multidrug-resistant bacterial infections are a major threat to public health, demanding new therapeutic solutions. Bacteriophages offer a promising alternative to antibiotics^1^, but their clinical use is constrained by a fundamental limitation: the narrow host range of individual phages, particularly against capsulated bacteria such as *Acinetobacter baumannii* and *Klebsiella pneumoniae*.^12,13,18,19,20^ Current approaches rely on assembling large libraries of genetically diverse phages^14^, each requiring separate process development and regulatory approval.^10,11^ This places a significant, often prohibitive, burden on phage therapy development. In principle, synthetic biology provides a way forward by enabling rational modification of phages to alter their host range without the need to redevelop entire therapeutic agents from scratch. Specifically, standardized therapeutic phage scaffolds could be configured with diverse natural RBPs that attach to the scaffold through their compatible adapter domain.^23,25,28,29^

In this study, our analyses establish that N-terminal adapter domains of RBPs form large clusters (Figure 2A). TAILOR reliably identifies these domains, with predictions consistent across benchmarks (Figure 1D), 3D structural models, and experimental swap data. While adapters are often conserved, particularly in Autographivirales phages, their associated enzymatic domains are highly diverse, generating a wide range of capsule specificities (Figures 3-4). This diversity scales with adapter cluster size, suggesting extensive natural diversity (Figure 3B). Linking these clusters to global pathogen populations shows that a relatively small number of adapter clusters can cover a large fraction of carbapenem-resistant *A. baumannii* and *K. pneumoniae* (Figure 4). Together, these results provide a foundation for the systematic configuration of therapeutic phages with clinically relevant host recognition RBPs.

Several limitations of our approach must be considered. First, although RBP swaps can confer complete host-range transfers in multiple cases^22,23,24,25,26,27^, RBPs are not the sole determinants of phage infectivity. Bacteria possess a variety of defense systems — such as CRISPR-Cas, restriction-modification systems, and abortive infection mechanisms — that can inhibit infection even after receptor recognition.^43^ These defense systems remain poorly annotated across clinical isolates and may substantially impact phage efficacy in vivo. Incorporating bacterial defense and phage anti-defense mechanisms into our framework represents an important next step for designing more effective phages. Second, the assumption of uniform phage susceptibility among strains sharing the same genomically predicted capsule type can be an oversimplification. Recent work on *A. baumannii* showed that phage susceptibility can be homogeneous across multiple capsule types despite the tested isolates originating from different world regions, whereas for other capsule types, strains with identical capsule type predictions differ markedly in phage adsorption and susceptibility.^14^ For *K. pneumoniae*, such intra-type heterogeneity remains unexplored due to the lack of systematic phage susceptibility measurements on strain collections with identical capsule types, introducing uncertainty into the host range of RBP. Third, while our adapter clustering provides a strong foundation for engineering, experimental validation of putative compatibility groups remains limited. Nearly all documented RBP swaps involve Autographivirales phages, which is explained by the unique properties of their adapter clusters, making them particularly promising candidates for RBP exchange. Further systematic experimental efforts to validate these compatibility predictions will be critical for confirming their utility in constructing therapeutic phage collections with nearly identical genomes. Finally, we substantially underestimate the actual diversity of specificities in large adapter clusters, as many predicted depolymerases lack specificity assignments due to the presence of multiple depolymerases in the corresponding phage genomes or a lack of experimental data. Therefore, experiments characterizing RBP specificities can quickly improve the coverage of high-priority pathogen populations.

## Conclusions

We established a computational framework to identify host recognition protein repertoires for plug-and-play configuration of therapeutic phages to cover highly diverse pathogen populations. We found that adapters — especially in Autographivirales phages — are widely conserved structurally while they are associated with highly diverse enzymatic domains. Due to this diversity in host specificities, a limited number of adapter clusters can collectively target a substantial fraction of carbapenem-resistant *A. baumannii* and *K. pneumoniae* populations. Future work should focus on expanding RBP annotations, validating adapter compatibility systematically, and incorporating bacterial defense systems and capsule heterogeneity into programmable phage design. Together, these efforts will be crucial for translating modular phage engineering from a few conceptual showcases into robust design principles.

## Methods

### Data acquisition and preprocessing

We retrieved all publicly available *A. baumannii* (30,827 as of 24 January 2025) and *K. pneumoniae* (55,742 as of 31 December 2023) genomes from NCBI. We confirmed the species designation of each genome using Kraken 2^48^. For *A. baumannii,* we performed capsule typing using Kaptive^19,49^ and looked for carbapenem resistance genes by predicting open reading frames (ORFs) using Prodigal v2.6.3.^50^ Then, we identified resistance genes based on sequence similarity to literature-curated resistance genes compiled in the ResFinder database v2.0.0.^51^ For *K. pneumoniae* we performed both capsule typing and resistance gene detection using Kleborate v2.4.1.^52^ For both species, we retained only those genomes that met the following criteria: i) confirmed species designation, ii) reliable capsule prediction (at least ‘Good’ with Kaptive2 and Kleborate, ‘Typeable’ with Kaptive3), iii) presence of at least one carbapenem resistance determinant, iv) available metadata on time and location of sample collection, and v) human-relatedness. In addition, we restricted our analysis to isolates collected between 2018 and 2023. After filtering, we obtained final sets of 16,031 and 10,810 genomes for *A. baumanni*i and *K. pneumoniae*, respectively. Since the filtered set of isolates was likely biased due to uneven representation of regions and time periods, we corrected for this by stratifying the data by geographical region and time, and then by population. Specifically, we applied the following subsampling procedure: First, for isolates with known city information, a maximum of one sample per city per week was kept, and isolates with only country information were limited to a maximum of two samples per country per week. Next, each country was limited to a maximum of one sample per million inhabitants. Ultimately, 2,946 *A. baumannii* and 2,959 *K. pneumoniae* isolates remained after downsampling (Table S20).

We collected a list of Acinatobacter and Klebsiella phage genome IDs from the International Committee on Taxonomy of Viruses (ICTV), and INPHARED^44,45^ (as of October 2024) and downloaded these genomes from NCBI. We attached taxonomy information using the NCBI taxonomy database, through the R package taxizedb v0.3.2^47^, and host information using data from the Virus-Host DB database (https://www.genome.jp/virushostdb/). From the annotated genomes, we selected proteins whose annotation notes contained the following strings: *tail fiber, tail fibre, tail spike, spike, fibre, fiber* and excluded those that contained: *appendage, assembly, connect, detail, head, base, attachment, socket, chaperon, joining, knob, measur, neck, needle, sheath, tip, tube, tubular*.

In the case of the phages from our collection^14^, we used Phold^53^ to identify tail fibers and spikes from phage genomes. This preprocessing step yielded 2,315,858 protein sequences, which were reduced to 1,109,499 unique sequences after deduplication at 97% pairwise identity. Of these, 2,313 were identified as suitable RBPs for our analysis, originating from 1,270 phage genomes (Table S1). Before proceeding with downstream analysis, we deduplicated all RBPs at a 97% amino acid identity cutoff and used the representative sequences to identify N-terminal adapter domains and cluster them.

### Detection of N-terminal adapter domains

The RBPs of phages typically contain a conserved N-terminal adapter region, which anchors the RBP to the phage particle^32^ (Figure 1A). Throughout this paper, the term ‘adapter’ refers specifically to the N-terminal region of the RBP, which differs from the usage of ‘adaptor’ by others for the gp11 protein^54^, while the terms receptor binding protein (RBP), tail protein, and tail spike are used interchangeably. To detect these conserved adapter regions, we compared all unique amino acid sequences of the RBPs using pairwise global alignment with the BLOSUM-80 substitution matrix, implemented in the R package pwalign v1.4.0.^55^ When two RBPs have similar N-terminal adapter regions but differ elsewhere, their alignments typically show many positive substitution scores in the N-terminal region, followed by predominantly negative substitution scores afterwards.

The end position of the adapter region (*i.e.*, the breakpoint) was identified using a two-step approach: First, we calculated the percentage identity across the initial 5% of the overall alignment length. This value was used as the center for an xbar.one control chart (one-at-a-time data of a continuous process variable) implemented in the R package qcc v2.^56^ Violations were used to detect similarity regions in the alignment. Mean substitution scores and percentage identity values were calculated for each such region. We then searched for a conserved N-terminal region that starts in the first quarter of the alignment length and has at least 40% sequence identity. If such a region was found, the original alignment was carried forward for fine-tuned breakpoint detection. In this case, we used the R package breakpoint v1.2^57^with the cross-entropy method to detect breakpoints between alignment sections with different similarity scores. As with the xbar.one approach, we used this method to detect similarity regions within the alignment. Similarly, mean substitution scores and percentage identity values were calculated for each region, and then consecutive conserved regions were merged. The end position of the shared N-terminal adapter domain was calculated as the end of the first conserved region, which begins in the first quarter of the alignment and has at least 40% sequence identity. We used this two-step approach because the xbar.one control chart was more effective at detecting the presence of a shared conserved N-terminal domain, whereas the cross-entropy method more accurately identified the end position of the domain.

### Clustering the RBPs based on adapter domain similarity

Once N-terminal adapter domain candidates were annotated using pairwise global alignments, we assigned each RBP to adapter clusters based on adapter domain sequence similarity. To this end, we calculated a Euclidean distance matrix based on adapter sequence similarities and then performed average-linkage hierarchical clustering using base R functions dist() and hclust(). To determine the optimal threshold for separating the clusters, we tested multiple values and evaluated the resulting clusters based on their biological relevance.

For this evaluation, we developed simplified metrics for homogeneity and completeness, inspired by previous studies.^58,59^ A 75% identity threshold was used to define a shared adapter between a pair of RBPs; that is, two RBPs were considered to share an adapter if their N-terminal domains showed at least 75% sequence identity. Homogeneity measures the similarity of samples within a cluster and was assessed here as the percentage of RBP pairs that shared adapters within a cluster, relative to the total number of pairs in that cluster. Note that if every pair of RBPs in a cluster shared the same adapter, the cluster was complete, with a homogeneity score of 1. Clusters with a homogeneity score below 0.1 were subjected to a second round of clustering, producing subclusters indicated by a trailing number (*e.g.*, ACL 2.1, where ACL stands for Adapter Cluster, while 2.1 indicates that it is a subcluster derived from cluster 2). Completeness measures whether all proteins that share N-terminal domains with members of a cluster are also included within that cluster. If all proteins that share the same N-terminal domain with cluster members are contained within the cluster (*i.e.*, cluster members don’t share N-terminal domains with any proteins outside the cluster), that cluster will have a completeness score of 1. We evaluated all possible clustering scenarios in a stepwise manner: we used the base R function cutree() to cut the hierarchical clusters into 1, 2, …, N clusters, where N was the overall number of RBPs, calculated homogeneity and completeness scores for each cluster, and finally we calculated a single weighted sum of scores for each scenario, in which individual homogeneity and completeness scores were weighed by cluster size. Ultimately, the optimal number of clusters was determined by maximizing this score, thereby grouping the RBPs into relatively few homogeneous and complete clusters.

The length of matching N-terminal sequences depends on the pair being compared; thus, there is no single adapter length for an RBP or cluster, but rather a range of values. However, for downstream analyses, we assigned a single adapter length to each cluster. To this end, we examined all pairwise comparisons within each cluster and calculated the median length of the identified adapter sequences.

We evaluated the quality of each adapter cluster. For each pair of sequences within a cluster, we retrieved the percentage identity of their shared adapter sequence. We considered an adapter cluster of good quality if the median percentage identity of these shared adapters was at least 40% and their standard deviation was below 30%. We defined the top adapter clusters as those of high quality and containing at least 30 RBPs.

### Prediction of enzymatic domains

In a complementary approach, we used InterProScan^60^ and Phyre2^61^ tools to identify enzymatic domains in RBPs. InterProScan predicts protein domains and functional sites by comparing protein sequences to several databases (*e.g.*, Pfam, SMART, TIGRFAMs), while Phyre2 predicts protein structure using homology modelling. To ensure that only reliable domain predictions are retained, for InterProScan, we kept domain predictions with an e-value ≤ 10. For Phyre2, we considered only matches with a confidence score of at least 0.4 and a minimum domain length of 60 amino acids. With both tools, we only retained those results that contained at least one of the following keywords, irrespective of case: *hydrolase, lyase, hyaluronidase, sialidase, rhamnosidase, levanase, dextranase, xylanase, depolymerase, glycos,* and *endosialidase*. Then, we excluded any enzymatic domain matches that overlapped with our adapter domain predictions by more than 25% of the length of the shorter amino acid sequence. Finally, from the remaining InterProScan and Phyre2 hits, we selected the longest match from each. If there were both an Interpro and a Phyre2 hit corresponding to the RBP, we prioritized Phyre2 as it gave us more realistic domain lengths and more informative descriptions. If there were neither Interpro nor Phyre2 predictions passing our requirements, we concluded that the RBP had no enzymatic domains. We found that the overwhelming majority of enzymatic domains followed the adapter region and showed no or minimal overlaps with it (Supplementary File 2).

### Assigning capsule specificities to RBPs

To assign capsule specificities to RBPs in our database, we conducted a literature review to collect publicly available capsule specificity data and then matched known specificities to RBPs in our database based on sequence similarity. First, we looked for articles about bacteriophages that infect *Acinetobacter* or *Klebsiella* hosts with known K types. If an amino acid sequence was missing from the publication, we attempted to retrieve the sequence from NCBI using the R package rentrez v1.2.4.^62^ For each protein, we determined the associated host organism through a series of queries that again retrieved data from NCBI, using functions from the R packages rentrez v1.2.4, webseq v0.1.0^46^, and taxizedb v0.3.2.^47^ We confirmed that all of these proteins were associated either with Acinetobacter or Klebsiella hosts. We only proceeded with proteins where we could find the amino acid sequence. Furthermore, in the case of proteins from the publication by Beamud and coworkers^16^, we only proceeded with proteins that belonged to RBD clusters with at least 50% True Positive Rate (TPR). We also added 6 Acinetobacter RBPs to this list from our previous study.^14^ These 6 proteins were the only RBPs in their phages with an annotated depolymerase, and so we assumed that these RBPs can be associated with the K type of the host that they successfully infect. Overall, we found 247 proteins in our literature review, out of which we progressed 123 (Table S13). Next, we grouped the entries by amino acid sequence and pooled specificity values. Grouping resulted in a non-redundant set of 94 amino acid sequences, out of which 31 and 63 were associated with Acinetobacter or Klebsiella hosts, respectively (never both). To harmonise specificity values, we used the K locus reference databases from Kaptive v3.1.0^49^ and coerced to K types everywhere to remove ambiguity. However, for K locus types where Kaptive reported unknown CPS types, we kept the K locus type, for simplicity, *e.g.*, “KL106” instead of “Unknown (KL106)”. Next, we used Protein-Protein BLAST (blastp) v2.13.0+ to search our database for RBP sequences. We retained only hits with at least 95% identity, 90% coverage, and matching host organisms (*e.g.*, for an RBP from an Acinetobacter phage, we only accepted RBP hits that were also from an Acinetobacter phage). Finally, when there were multiple valid hits for an RBP, then matched specificities were again pooled, and the resulting list of unique specificities was assigned to the RBP (Table S1).

### Predicting the structure of RBPs

To make the most effective use of our resources, we decided to proceed with a subset of RBPs that allows us to work with a representative sample of the dataset-capturing as many enzymatic clusters as possible, minimizing sequence redundancy, and softly prioritizing proteins from phages available for experimental validation.

AlphaFold2-multimer v2.3.1^63^ was employed to predict the structures of RBPs, using an Nvidia A100 GPU with 5 seeds and no relaxation applied. The resulting models were ranked based on two confidence scores: the pLDDT (predicted local-distance difference test) score and the interface-predicted TM-score (iPTM), with the top-scoring model being selected for further classification. pLDDT is a per-residue confidence score generated by AlphaFold, which evaluates the confidence in the local structure of individual residues, while the iPTM score assesses the confidence in the predicted interfaces between protein chains. By selecting the model with the highest combined score, we ensured that both local structures and inter-chain interfaces are predicted with high confidence. Structural similarity among the models was assessed using US-align^64^, which was used to superimpose and score the structures based on their TM-score. Subsequently, k-means clustering -an iterative process to minimize the sum of distances between the data points and their cluster centroids-was applied to group the models based on their structural similarities. The N- and C-terminal regions of each structure were manually isolated using ChimeraX v1.6.1, after which the same clustering procedure was performed on these regions to identify patterns specific to the termini.

### Visual inspection of 3D models

To identify the end of adapters in our predicted 3D structures, we considered the end of the alpha helices, just before the beginning of the depolymerase, to be the end of the adapter structure. When there was no traditional adapter structure (in the case of very short adapters dubbed heptamers by Latka and coworkers^14^, we used the beginning of the depolymerase, after the seemingly disordered amino acids at the N-terminal, as the delineation point. Out of 105 RBPs with predicted 3D structures 57 were evaluated by this visual inspection (Table S7).

## Availability of data

TAILOR is an R package freely available on GitHub (https://github.com/sbthandras/tailor). The datasets supporting the conclusions of this article are included within the article and its additional files.

## Acknowledgements

This work was supported by the National Research, Development and Innovation Fund ADVANCED 149516 grant (BK), the 2024-1.2.2-ERA_NET-2024-00004 grant (BK and BP), the National Laboratory of Biotechnology grant 2022-2.1.1-NL-2022-00008 (BK), the National Laboratory for Health Security grant RRF-2.3.1-21-2022-00006 (BP), the HUN-REN TKCS-2024/66 grant (EA and BP), the Ministry of Culture and Innovation of Hungary grant KDP-2023-C2285907, financed under the National Research, Development and Innovation Fund 2023-2.1.2-KDP-2023-00002 funding scheme (TS); the European Union’s Horizon 2020 Research and Innovation Programme no. 739593 (BK and BP). We thank Tibor Páli, Koto Teruaki, and Hugo Oliveira for useful discussions.

## Contributions

AA, TS, BK and GA developed the conceptual design of the project. AA, TS and GA performed data acquisition and analyses. AA, TS, OM, EA and BK interpreted results. AA, TS, OM and EA generated figures. HHM was responsible for literature acquisition and reference management. AA, TS, OM, EA, BK and BP wrote and revised the manuscript. BK, BP and EA supervised data generation and analyses. VK performed the RBP structure predictions and NT supervised structure predictions. All authors provided feedback and approved the final manuscript.

## Corresponding author

Correspondence to Bálint Kintses

## Ethics approval and consent to participate

Not applicable.

## Consent for publication

Not applicable.

## Competing interests

Balint Kintses is the founder of SplitPhage Solutions Ltd. and a shareholder in BRC-Bio Ltd., companies operating in the field of biotechnology. These companies had no role in the design, execution, interpretation, or funding of this study.

## Supplementary Figures

**Figure S1.**
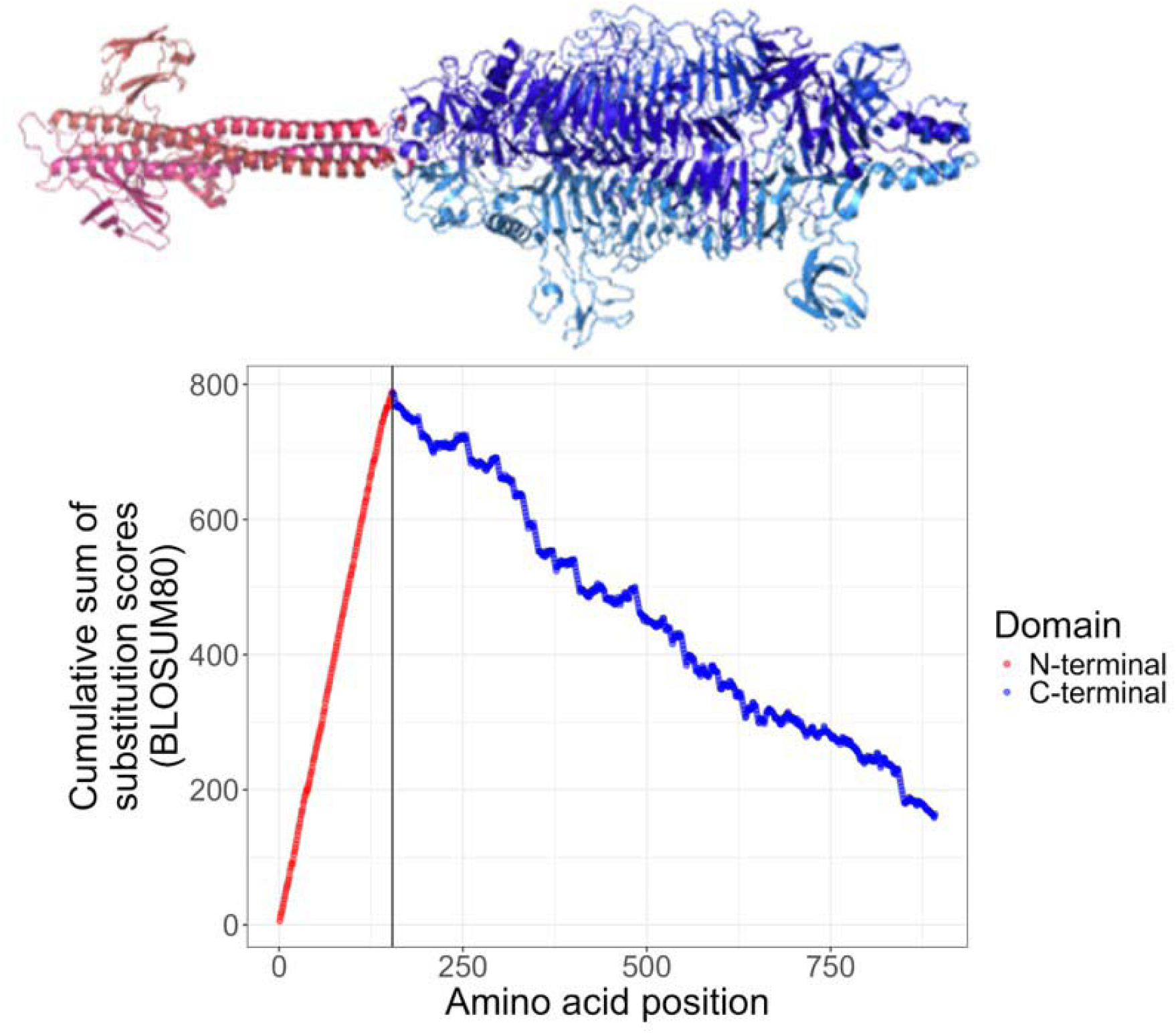
Predicting the N-terminal adapter domain by defining the boundary (vertical black line) between the conserved N-terminal (red) and the variable C-terminal regions of RBP. The boundary was defined based on a drop in the cumulative substitution score across all unique RBP sequences, using pairwise global alignments (see Methods). The figure illustrates a specific example derived from the alignment of RBP sequences from *Acinetobacter* phage vB_AbaP_IME546 (BDJVYHBB_CDS_0054) and *Friunavirus* AS11 (PCQZKNGA_CDS_0051). The x-axis indicates amino acid positions, numbered from the N-terminal end of the RBP (see Table S1 and S2).

**Figure S2.**
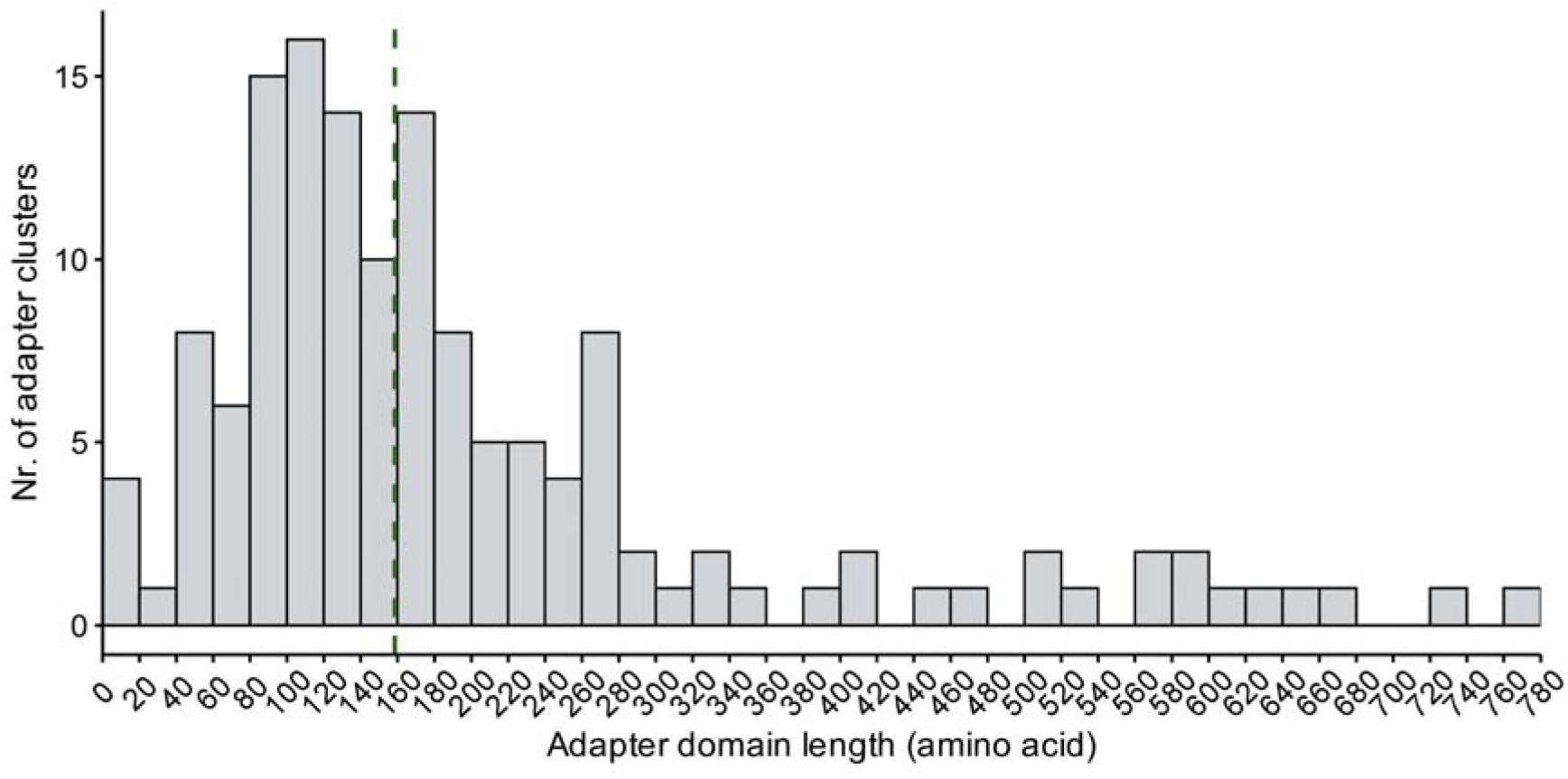
Most adapter domains range in length from 100 to 240 amino acids, with a median length of 158 (indicated by the green vertical line). Length was calculated only for adapter domains belonging to the 172 non-singleton clusters. Notably, 18 clusters contain predicted adapter domains ranging from 100 to 120 amino acids in length. For more information on how adapter lengths were calculated (see Methods and Table S5). Two outlier values (1835 and 1875) were excluded from the figure to improve visual clarity.

**Figure S3.**
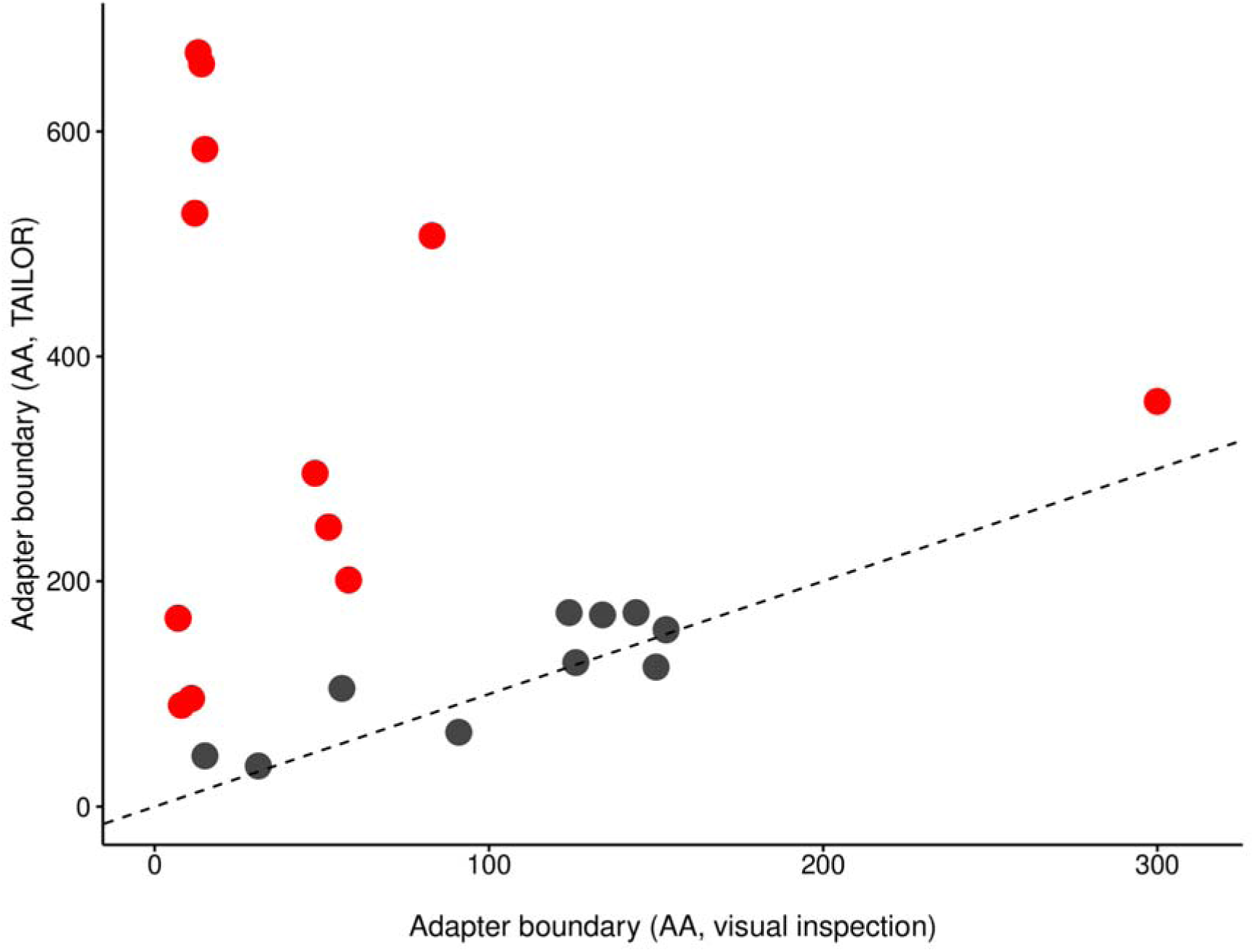
Predicted adapter domain lengths from TAILOR are similar to domain lengths inferred by visual inspection of the 3D structure. While TAILOR tends to predict somewhat longer adapters than those assumed by visual inspection, adapter domain lengths from TAILOR and from visual inspection correlate well (Pearson correlation coefficient 0.868, for the 10 black points). Adapters shorter than 15 amino acids or predicted to be longer than 200 amino acids, where TAILOR often could not detect the right sequence are highlighted in red. The dashed line marks identity (Table S7). The figure is based on Table S3 and only those cases are shown where the median adapter length predicted by TAILOR was 200 amino acids or less (see Table S7).

**Figure S4.**
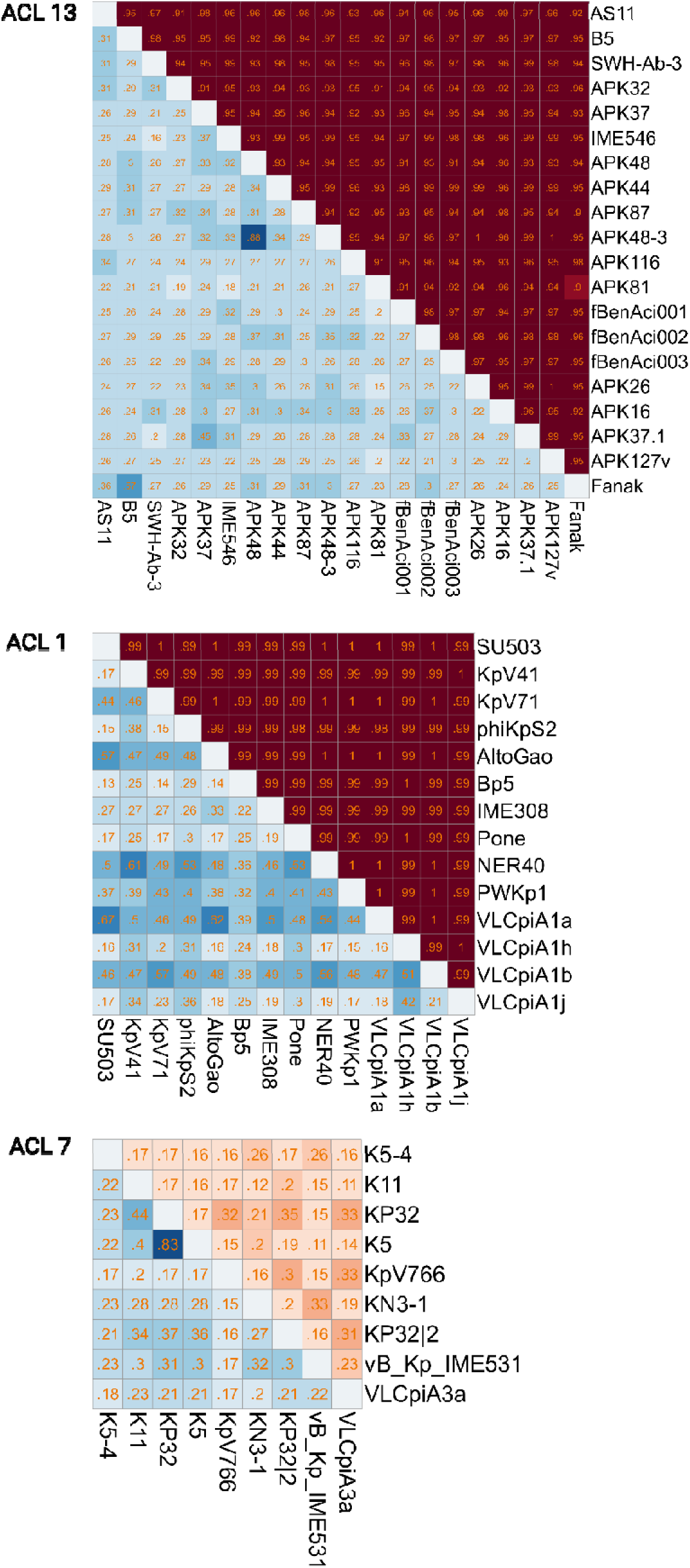
Heatmap showing pairwise TM-score-based structural similarities of N-terminal adapter domains (upper right part, shades of red) and C-terminal domains (lower left part, shades of blue) of phages’ RBPs within the ACL 13, ACL 1 and ACL 7 clusters. Within ACL 7 the structural similarity of adapter domains is not significantly higher than that of the C-terminal domains (p-value = 0.9993 from one-sided Wilcoxon test), as the 20–36 amino acid-long adapters from this cluster do not exhibit a highly conserved structure. TM score values are indicated in the boxes as abbreviated decimal numbers. The names of phages to which the RBPs belong are indicated on the x and y axes (Table S9).

**Figure S5.**
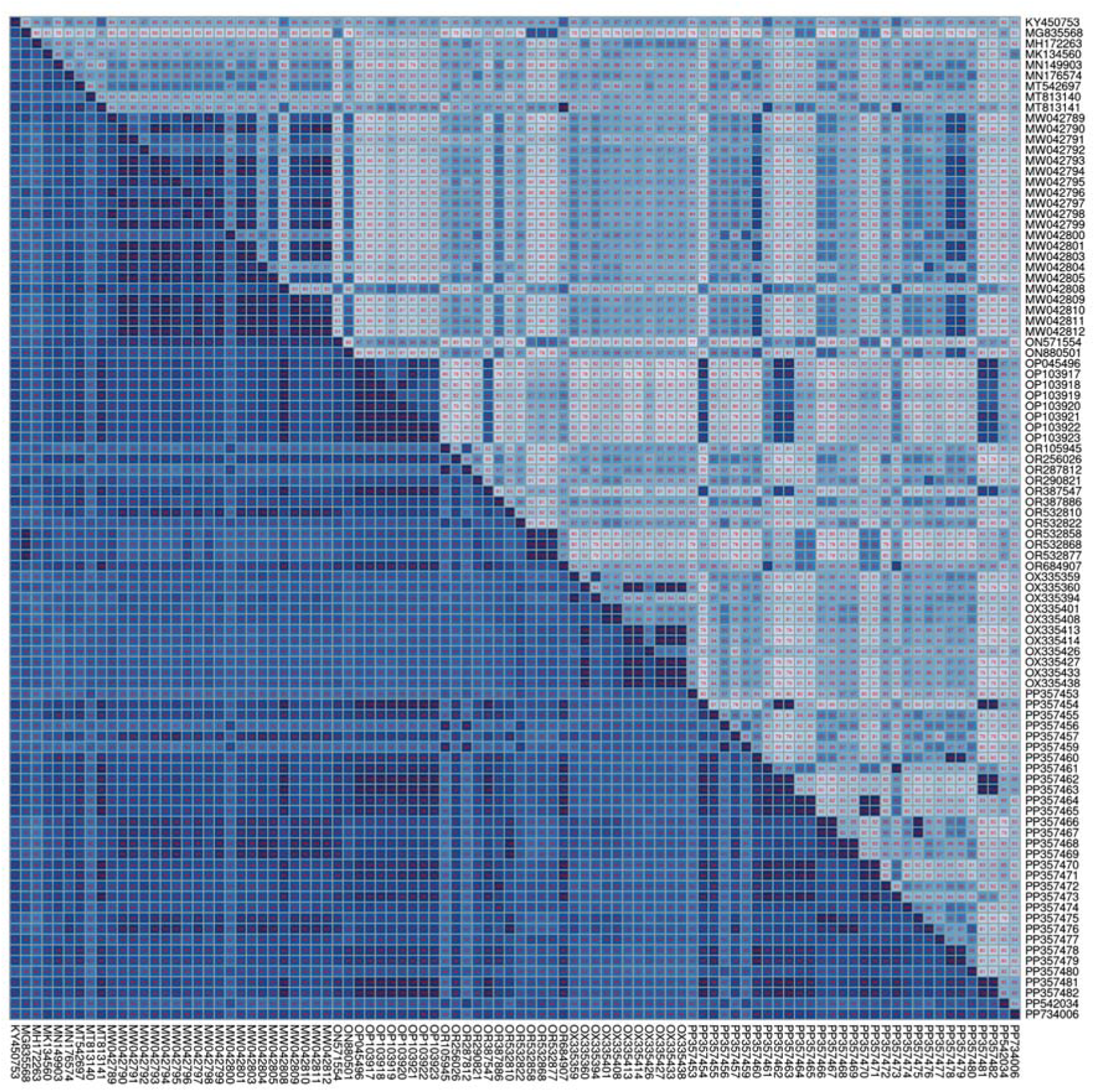
94 distinct *Klebsiella* vOTUs share an RBP with very high sequence identity along the entire length, including the adapter that belongs to ACL 6. Pairwise average nucleotide identity (ANI) values were calculated using BLASTN. Upper triangle: ANI across *Klebsiella* vOTUs (N = 94). Lower triangle: ANI across RBP sequences. Values represent length-weighted percent identity with correction for overlapping alignments (Table S23).

